# Probenecid Inhibits SARS-CoV-2 Replication *In Vivo* and *In Vitro*

**DOI:** 10.1101/2021.05.21.445119

**Authors:** Jackelyn Murray, Robert J. Hogan, David E. Martin, Kathy Blahunka, Fred Sancillo, Rajiv Balyan, Mark Lovern, Richard Still, Ralph A. Tripp

## Abstract

Effective vaccines are slowing the COVID-19 pandemic, but SARS-CoV-2 will likely remain an issue in the future making it important to have therapeutics to treat patients. There are few options for treating patients with COVID-19. We show probenecid potently blocks SARS-CoV-2 replication in mammalian cells and virus replication in a hamster model. Furthermore, we demonstrate that plasma concentrations up to 50-fold higher than the protein binding adjusted IC_90_ value are achievable for 24h following a single oral dose. These data support the potential clinical utility of probenecid to control SARS-CoV-2 infection in humans.

## Introduction

Severe acute respiratory syndrome coronavirus 2 (SARS-CoV-2) causes coronavirus disease 2019 (COVID-19). In late 2019, the world became aware that SARS-CoV-2 had emerged as a pandemic confirmed by the World Health Organization (WHO) in March 2020. As of late April 2021, there are >141 million COVID-19 cases with >3 million deaths worldwide according to the COVID-19 Data Repository by the Center for Systems Science and Engineering (CSSE) at Johns Hopkins University. The SARS-CoV-2 pandemic underscores the urgent need for therapeutics for treatment or prevention as the current armamentarium is comprised of three vaccines^1–3^, palliative support, and one antiviral drug, i.e. remdesivir. Remdesivir is a nucleoside analog granted emergency use authorization (EUA) by the FDA based on the demonstration of a decreased time to recovery in patients hospitalized for severe COVID-19^4^. The use of remdesivir is restricted to treating hospitalized, COVID-19 patients via intravenous administration. Vaccines and small molecule therapeutics targeting viruses are the mainstays of antiviral approaches, but complementary approaches targeting host cell genes/processes are needed to reduce the propensity for the development of drug resistance and because targeting host cell pathways can confer broad-spectrum antiviral activity^5,6^.

Identifying host cell targets crucial to virus replication is facilitated by high-throughput screening (HTS) methods including the use of RNA interference (RNAi) methodologies^7^. This approach has been used to identify and validate molecular targets and pathways having previously unappreciated antiviral actions, particularly for drug repurposing^6,8^. The repurposing of existing drugs with a known clinical profile has significant advantages over drug discovery as it lowers the risk, time, and cost for development and entry into the clinic^9^. Using HTS to discover host genes required for viral replication, the organic anion transporter 3 (OAT3) gene was identified as a host gene that could be repurposed as a therapeutic candidate^10^.

OAT3 is expressed in the kidney, choroid plexus, vascular beds, and other peripheral organs including the lung^11^, and mediates the transmembrane transport of endogenous organic anions including urate and other substrates and certain antibiotics^12^. Probenecid {4-[(dipropyl-amino) sulfonyl] benzoic acid} is a commonly used therapeutic agent that inhibits OAT3^13^. Probenecid is a gout treatment, and is a favorable candidate for antiviral drug repurposing, as it is readily commercially available with favorable pharmacokinetics and has a benign clinical safety profile^14^. Probenecid prophylaxis and treatment have been shown to reduce influenza virus replication *in vitro* in mice^10^. The half-maximal inhibitory concentration (IC_50_) for treatment of A549 human type II respiratory epithelial (A549) cells infected with A/WSN/33 (H1N1) or A/New Caledonia/20/99 (H1N1) was 5 x 10^-4^ and 8 x 10^-5^ uM, respectively^10^.

We determined the IC_50_ for Vero E6 cells and normal human bronchoepithelial (NHBE) cells treated before and at the time of infection with SARS-CoV-2 and the B.1.1.7 variant. The results showed probenecid treatment blocked SARS-CoV-2 replication (plaque formation) in Vero E6 cells or NHBE cells treated with different concentrations of probenecid, i.e. 0.00001 - 100 uM. We also evaluated lung viral load in hamsters on days 0, 3, and 7 pi with SARS-CoV-2 using both a tissue culture infectious dose-50 (TCID_50_) assay and a virus plaque assay to determine the number of plaque-forming units (PFU). Hamsters were treated with probenecid 24h before infection (prophylaxis), or 48h postinfection (post-treatment) with doses of 2 mg/kg and 200 mg/kg. Hamsters treated with probenecid had dramatically reduced lung virus titers, i.e. a 5-log reduction of virus compared to the controls that were approximately 10^9^ logs of the virus. We performed a probenecid pharmacokinetic modeling and simulation study comparing 600 mg twice daily, 900 mg twice daily, or 1800 mg once daily. The model predicted that the plasma concentrations of probenecid would exceed the protein binding adjusted IC_90_ value at all time points throughout therapy. The doses evaluated by the PK model were below the maximum allowable FDA-approved dose of 2 grams/day and should have no substantial side effects. Together, these data strongly support the potential for probenecid to provide a robust antiviral response against SARS-CoV-2.

## Results

After performing RNAi screens in human lung type II epithelial (A549) cells that allowed us to determine and validate host genes required for influenza virus replication^6^, we chose to evaluate OAT3^10^. Transfection of A549 cells with siRNA targeting the SLC22A8 gene, OAT3, completely blocked influenza A/WSN/33 (H1N1) virus replication, and probenecid treatment reduced OAT3 mRNA and protein levels *in vitro* and BALB/c mice. As probenecid treatment *in vitro* or *in vivo* did not block influenza virus infection but did block virus titers as measured by plaque assay and hemagglutination assay, we determined the in vitro inhibitory effect on SARS-CoV-2 replication in Vero E6 cells, and NHBE cells (Figure 1). Vero E6 cells and NHBE cells were pretreated with differing probenecid concentrations and the effect on viral RNA load in the tissue culture supernatant was determined at 48h after infection by plaque assay. Probenecid treatment resulted in a dramatic decrease in SARS-CoV-2 replication by nearly 90% in Vero E6 cells and >60% in NHBE cells when compared to controls (Figure 1 A, 1B). The IC_50_ value for probenecid was shown to be 0.75 uM in Vero E6 cells and 0.0013 uM in NHBE cells. Viability was also assessed over the differing concentrations, demonstrating no cellular toxicity at the highest drug concentration (data not shown). Probenecid treatment was also effective at inhibiting a SARS-CoV-2 variant of concern (VOC), i.e. hCoV-19/USA/CA_CDC_5574/2020, B.1.1.7 (Supplemental Figure 2). This variant was isolated from a nasopharyngeal swab on December 29, 2020, and after sequencing it to the lineage B.1.1.7 (a.k.a. UK variant). This VOC may have increased transmissibility^16^.

**Figure 1.**
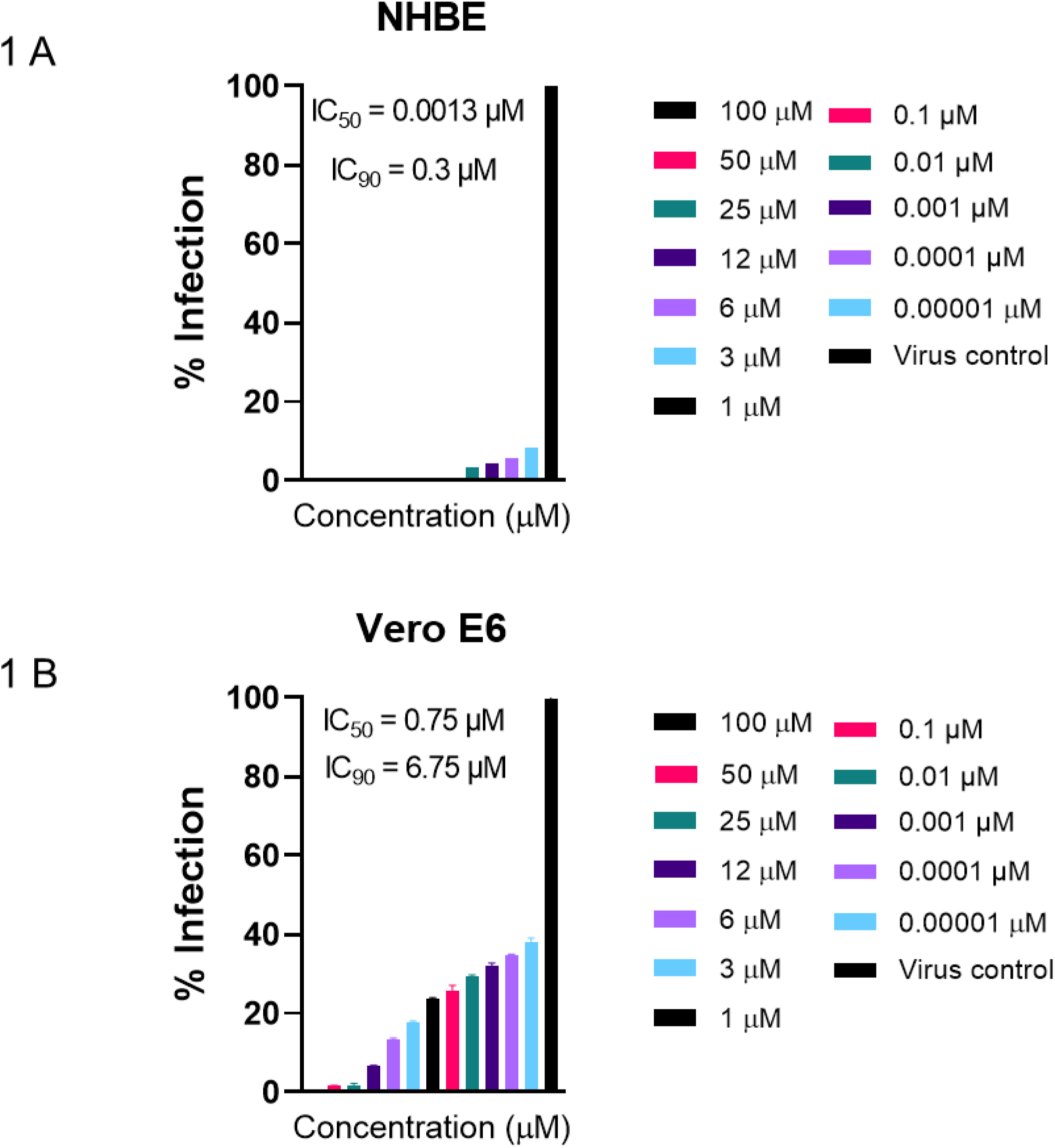
IC_50_ values for probenecid were calculated in NHBE cells and Vero E6 cells after treating with different concentrations of probenecid and detecting the virus culture supernatants by virus plaque assay. Probenecid treatment reduced SARS-CoV-2 replication by 90% in NHBE cells (A) or 60% in Vero E6 cells (B).

Having shown *in vitro* efficacy and determined the IC_50_ values, we next determined the efficacy of probenecid in the hamster model regarded as a preclinical model of SARS-CoV-2 disease with hamsters having self-limiting pneumonia^17,18^. We examined hamsters infected with SARS-CoV-2 and treated them with probenecid for either 24h before infection (prophylaxis) or 48h post-infection (post-treatment). Six dosing groups were tested: a) two prophylaxis groups treated with doses of 2 mg/kg or 200 mg/kg, b) two post-treatment groups treated with doses of 2 mg/kg or 200 mg/kg, and two control groups that were infected or not infected. Disease in hamsters following SARS-CoV-2 infection was transient peaking around day 3-4 post-infection with no clinical signs^17^. Consistent with this observation, no clinical symptoms or weight loss was evident in any group throughout the study (Supplementary Figure 1). Hamsters treated with probenecid had dramatically reduced lung virus titers (Figure 2), i.e. a 5-log reduction of virus compared to the controls that were approximately 10^9^ logs of the virus.

**Figure 2.**
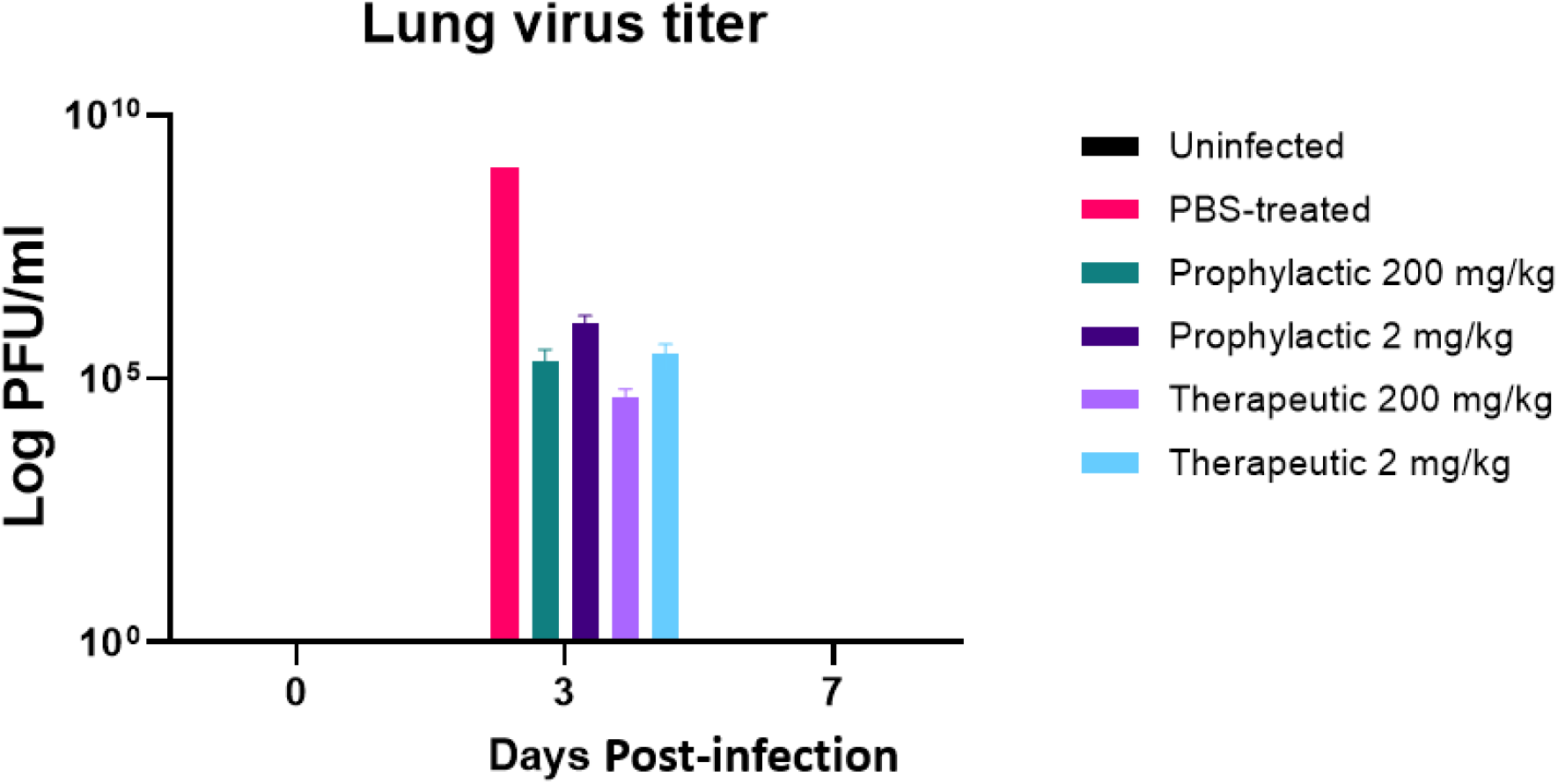
Hamsters were treated prophylactically or therapeutically with probenecid at 200 mg/kg or 2 mg/kg, intranasally infected with SARS-CoV-2, and the lungs viral titer determined at days 0, 3, and 7 pi. Control groups were uninfected or PBS-treated, infected hamsters. The viral titers were reduced by approximately 5-logs in the probenecid-treated groups compared to PBS-treated, infection control group.

Finally, a population pharmacokinetics (pop-PK) model was developed to characterize probenecid PK. It had a one-compartment structure with saturable elimination and first-order absorption. We performed simulations using the final pop-PK model to generate probenecid exposure profiles comparing 600 mg twice daily, 900 mg twice daily, or 1800 mg once daily administration (Figure 3). The doses examined are predicted to provide plasma concentrations exceeding the protein binding adjusted IC_90_ value at all time points. All doses were below the maximum allowable FDA-approved dose and are generally safe with no significant side effects.

**Figure 3.**
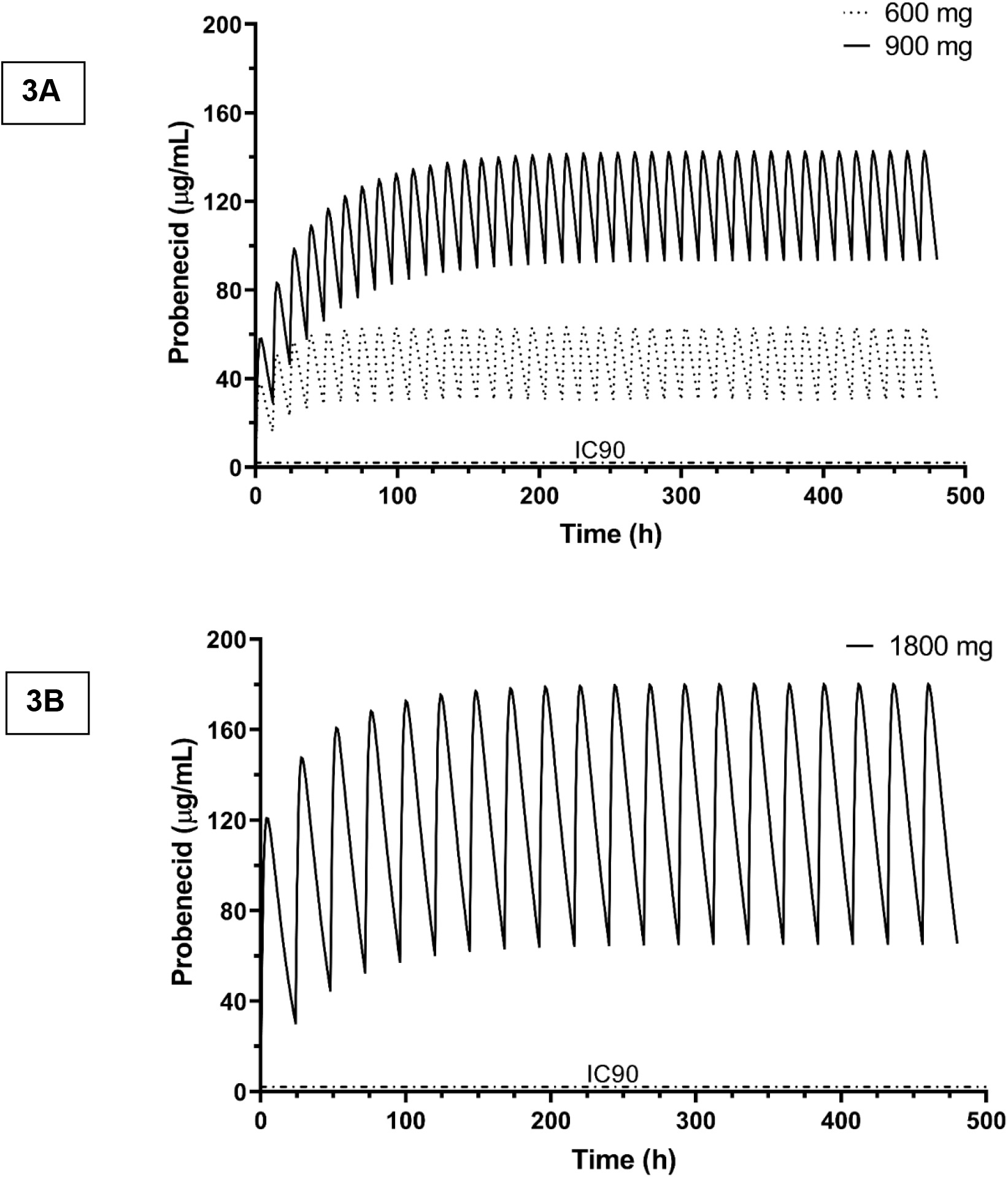
Simulated probenecid concentrations. A population pharmacokinetic model was used to generate probenecid exposure profiles comparing A) 600 mg twice daily, 900 mg twice daily, or B) 1800 mg once daily administration for 20 days. The IC_90_ level corrected for 95% protein binding is 2.08 μg/ml (shown as dashed line). All doses provided exposures well over the IC_90_ level at all time points.

## Discussion

The availability of small-molecule treatments for COVID-19 is limited. Currently, only remdesivir has received EUA approval for treatment of SARS-CoV-219 and it must be administered intravenously in hospitalized patients. There are several additional antivirals in development for the treatment of non-hospitalized patients. Table 1 compares the *in vitro* antiviral potency of remdesivir and several investigational agents against SARS-CoV-2. While all drugs show *in vitro* activity in the low micromolar to the submicromolar range, only probenecid can achieve the necessary plasma concentrations to produce an IQ value >2-fold. Importantly, probenecid has a long and benign safety record over 5 decades of use. Given *the in vitro* efficacy, the *in vivo* effect in hamsters, and the favorable pharmacokinetic results and benign safety profile, probenecid should be considered as a potential treatment option for COVID-19 patients.

**Table 1.** Comparative antiviral potency. Data comparing the *in vitro* antiviral potency of selected agents against SARS-CoV-2 and the potential inhibitory quotient (IQ). The IQ is used to express the ratio of plasma concentration at the trough to the protein binding adjusted IC_90_ value with higher values typically associated with greater clinical benefit.

| Compound | Company | Mechanism | Route | IC<sub>90</sub><br>(μM) | Cells | IQ* |
| --- | --- | --- | --- | --- | --- | --- |
| Remdesivir | Gilead | Nucleotide Analog | IV | 1.76 | Vero E6 | <2 |
| PF-07304814<sup>19</sup><br>(active: PF-00835231) | Pfizer | 3CL protease inhibitor | IV | 0.433 | HeLa-ACE2 | ~1 |
| PF-07321332# | Pfizer | 3CL Protease Inhibitor | PO | 0.16 | HAE* | <2 |
| AT-527<sup>20</sup> | Atea Pharmaceuticals | Nucleotide Analog | PO | 0.47 | HAE* | <2 |
| Molnupiravir<sup>21</sup> | Ridgeback BioTherapeutics | Nucleoside Analog | PO | 2.8 | HAE* | <2 |
| TD213 | TrippBio | OAT3 inhibitor | PO | 0.364 |  | ~50 |

Probenecid has inhibitory effects on RNA viruses specifically influenza^10^ and respiratory syncytial virus (RSV) (data not shown). Probenecid increases uric acid excretion in the urine and has been safely used to treat gout and hyperuricemia^22,23^, and has minimal adverse effects but may cause mild symptoms such as nausea, or loss of appetite^14^. Probenecid has several pharmacological targets including blocking pannexins that underlie transmembrane channel function^24,25^ and decreasing ACE2 expression^26,27^. The major advantages of probenecid are that it is an FDA-approved therapeutic drug that has been on the market for >50 years, it can be administered orally with favorable pharmacokinetics, it operates at the host cell level, is refractory to viral mutation, and has the potential to treat multiple other viruses. Further, our data suggest that initiation of posttreatment following infection in mammalian cells, hamsters, and humans results in infection remarkably reduced SARS-CoV-2 replication, particularly in the lung target organ.

Probenecid treatment will likely have the benefit of inhibiting SARS-CoV-2 variants as we show that it is effective against the VOC, lineage B.1.1.7. This is not unexpected, as targeting host processes essential for viral replication such as OAT3 would be expected to be universal. Among the host targets that have been identified as potential targets for inhibiting virus replication, OAT3 blockade will not likely confer any mechanism-based untoward effects for humans since humans with reduced OAT3 function are healthy^28^, and pharmacologic blockade of OAT3 is safely tolerated in humans^29^. Other compounds that interact with OAT3 include the antiviral drugs oseltamivir phosphate (Tamiflu) and acyclovir^10^, as well as angiotensin II receptor blockers^30^, however, their interaction is weak and their pharmacological actions confer safety and tolerability limitations that preclude their use in drug repurposing.

## Methods

### Biosafety

SARS-CoV-2 and variant studies were performed in BSL3 that was approved with relevant ethical regulations for animal testing and research. The hamster study received ethical approval from the University of Georgia IACUC committee and was performed by certified staff in an Association for Assessment and Accreditation of Laboratory Animal Care International-accredited facility. Work followed the institution’s guidelines for animal use, the guidelines and basic principles in the NIH Guide for the Care and Use of Laboratory Animals, the Animal Welfare Act, the United States Department of Agriculture, and the United States Public Health Service Policy on Humane Care and Use of Laboratory Animals.

### Animals

Syrian hamsters were group-housed in HEPA-filtered cage systems enriched with nesting material and were provided with commercial chow and water *ad libitum*. Animals were monitored at least twice daily throughout the study. Hamsters were intranasally infected with 10^4^ TCID_50_ SARS-CoV-2 (USA-WA1/2020; GenBank: MN985325.1). Hamsters were treated with probenecid 24h before infection (prophylaxis), or 48h post-infection (post-treatment) with doses of 2 mg/kg and 200 mg/kg. All groups were euthanized on day 3 or 7 pi, as day 3-4 pi has been reported as the peak of virus replication^31^.

### Hamster study design

Hamsters were divided into pre-infection or post-infection groups and as appropriate were intraperitoneally treated with probenecid (n = 6/group). Groups were treated with 200 mg/kg or 2 mg/kg of probenecid prophylactically at 24h preinfection, or prophylactically at 48h pi. Animal weights were collected once daily and animals were monitored twice daily for disease signs and progression. All procedures were performed on anesthetized animals. Lungs were collected on days 0, 3, and 7 pi for RT-PCR analysis.

### Virus and cells

SARS-CoV-2 isolate nCoV-WA1-2020 (MN985325.1) or the variant (hCoV-19/USA/CA_CDC_5574/2020, B.1.1.7; GenBank: MW422255.1) were received from BEI Resources managed under contract by American Type Culture Collection (ATCC). The viruses were propagated in Vero E6 cells and the cells maintained in high glucose DMEM supplemented with 10% fetal bovine serum, and 1 mM L-glutamine (tissue culture media, TCM).

### Virus load by RT-PCR

RNA was extracted from lungs using the QIAamp Viral RNA kit (Qiagen) according to the manufacturer’s instructions. Tissues were homogenized and RNA extracted using the RNeasy kit (Qiagen) according to the manufacturer’s instructions. qRT-PCR analysis of purified RNA was performed using the CDC EUA N1 probe assay and the CDC EUA human RNaseP internal control probe assay to validate sample RNA extraction in separate reactions.

### Virus titration

Virus isolation was performed by inoculating Vero E6 cells in a 96-well plate with a 1:10 dilution series of the virus, and one hour after inoculation of cells, the inoculum was removed and replaced with 0.2 ml TCM. Six days after inoculation, the cytopathogenic effect was scored and the TCID_50_ was calculated using the Reed-Muench method^32^.

### SARS-CoV-2 *In Vitro* Assay

Vero E6 cells (ATCC CRL-1586) were plated in 12-well plates at 5 x 10^5^ cells/well and incubated overnight at 37°C. Vero E6 cells were washed 1x with PBS and probenecid (Sigma) was added to the wells in DMEM media and incubated at 37°C. Each drug concentration was tested in triplicate and the experiment repeated >3 times independently. Following pre-treatment, DMEM media was discarded and cells were replenished with TCM containing probenecid and SARS-CoV-2. Cells were infected at an MOI = 0.01 for 4 days. Post-infection cells were fixed and stained to visualize plaques. Statistical analysis was by one-way ANOVA where p < 0.05. NHBE cells were sourced from LONZA (Walkersville, MD) from a non-smoking patient. The cells were seeded at 100,000 cells/T25 flask and incubated at 37°C. Once cells reached 70-80% confluency, they were dissociated using trypsin and plated using the conditions described above for culture with probenecid.

### Modeling and Simulation

Probenecid PK data was obtained from a published manuscript^33^. In this study, five adult healthy male subjects (20-39 years, 53-86 kg) were given oral 500, 1000, and 2000 mg probenecid at least one week apart and blood was collected until 48h. These data were fit in a nonlinear mixed-effects framework using Phoenix NLME software^34^. Parameter estimates for a PK model^33^ with saturating (Michaelis-Menten) elimination were used as initial estimates for the current analysis. The model was fitted using the first-order conditional estimation method with an extended least square (FOCE-ELS) algorithm. The base model testing included basic one and two-compartment structures with first-order absorption. Other absorption, lag time, and clearance models were also evaluated. A battery of diagnostic plots was employed to evaluate the adequacy of the goodness of fit for the PK model. The final popPK model was used to simulate exposures resulting from various dosing regimens of potential clinical interest (600 mg BID, 900 mg BID, 1800 mg QD).

## Acknowledgments

The authors would like to thank Phil Young and Billy Meadow for their advice. This work was funded in part by the Georgia Research Alliance and SpinUp Campus.

## Author contributions

J.M., R.H., D.M., K.B., and R.T. contributed to the design, data analysis; J.M., D.M., R.T. contributed to the writing of the manuscript; D.M., R.B., and M.L contributed to the probenecid modeling analysis; J.M., R.H. contributed experiment support; D.M. and R.T. contribute to the data analysis; all authors reviewed and contributed to the preparation of the final manuscript.

## Competing interests

D.E.M., K.B., F.S., R.S., and R.A.T have ownership in TrippBio, and all authors are knowledgeable of studies associated with TrippBio.

---

**Supplementary Figure 1.**
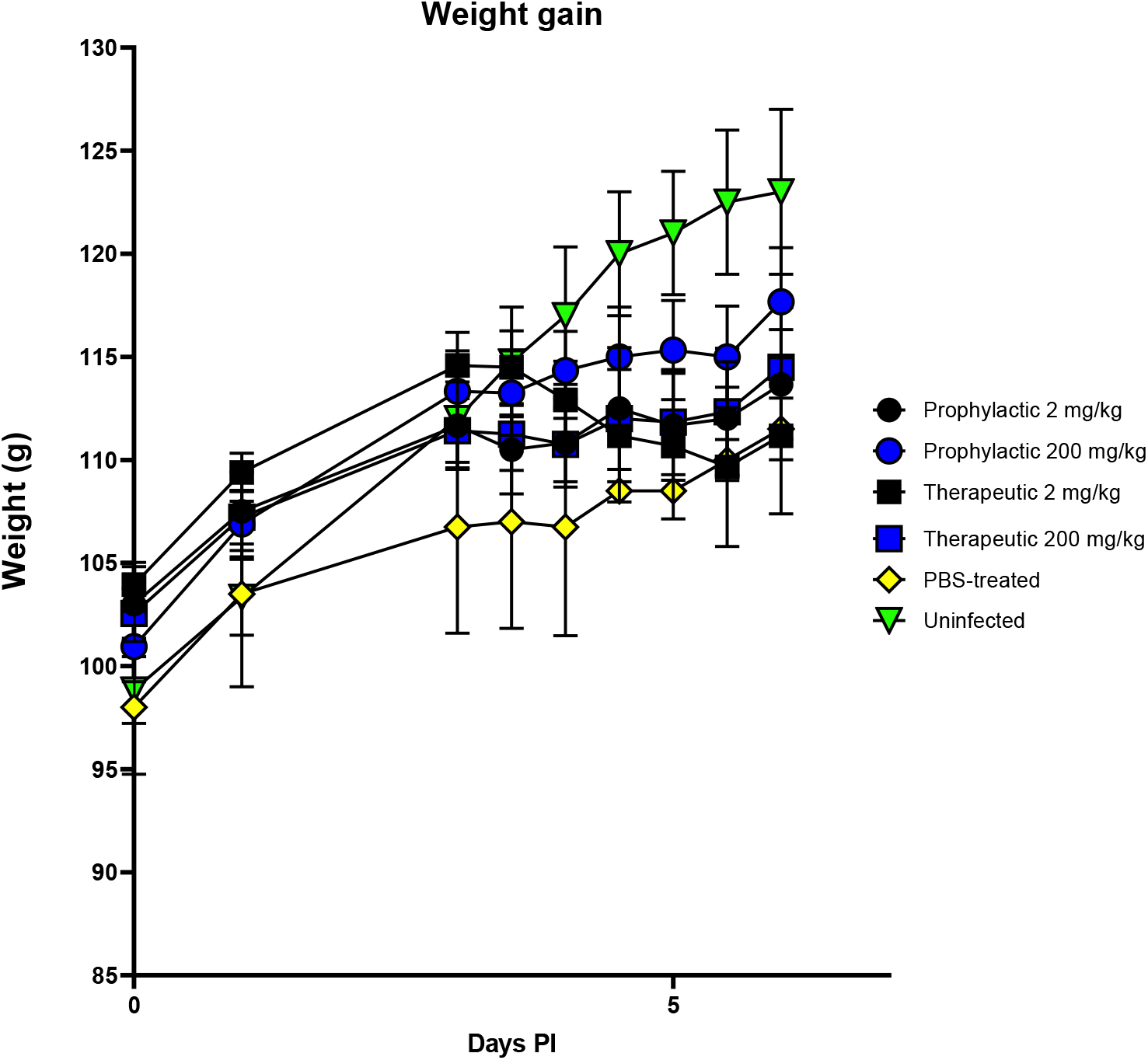
Hamsters were treated prophylactically or therapeutically with probenecid using 200 mg/kg or 2 mg/kg and intranasally infected with SARS-CoV-2. Uninfected hamsters or PBS-treated and infected hamsters were the controls. No weight loss was evident in any of the groups during this study.

**Supplementary Figure 2.**
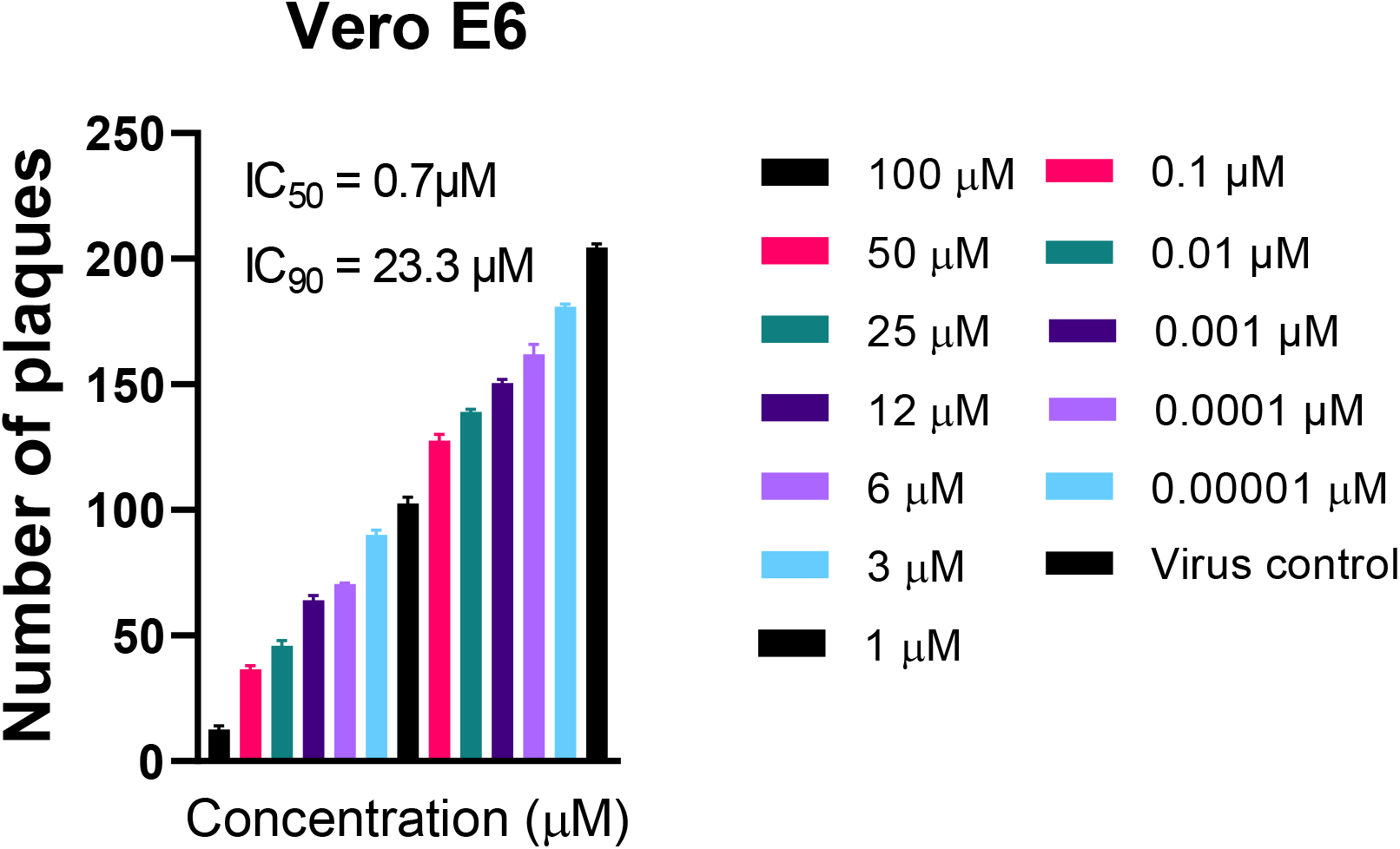
Probenecid treatment inhibited the replication of the SARS-CoV-2 variant of concern, hCoV-19/USA/CA_CDC_5574/2020, B.1.1.7. The IC_50_ value for probenecid was calculated in Vero E6 cells following treatment them probenecid concentration and determining the viral in culture supernatants by plaque assay.

## Notes

### Competing Interest Statement

D.E.M., K.B., F.S., R.S., and R.A.T have ownership in TrippBio, and all authors are knowledgeable of the studies connected to TrippBio.

